# How do age, available support, gender and attitudes affect the quality of data collected by young citizen scientists in an ecological research project?

**DOI:** 10.1101/2022.09.16.508269

**Authors:** Tuomas Aivelo

## Abstract

1: Citizen science is increasingly used to collect ecological data. Specifically, participation of school students in authentic research has been suggested as having a multitude of benefits from data collection to science education. Nevertheless, the overall quality and quantity of data concerns ecologists who are using data for further analysis.

2: I coordinate a citizen science project, the Helsinki Urban Rat Project where lower and upper secondary school students collect data on urban rat occurrence through the use of track plates that record rat footprints. Within the project, both the success and accuracy of participating young citizen scientists can be assessed by comparing their results by professional researchers’ results and due to the use of an additional questionnaire relate this to the background variables (i.e., age, gender, available support, attitudes and sensitivity to disgust).

3: I found out that, in contrast to results from previous studies, age was not a significant variable, but rather available support and voluntary participation with rewards increased data quality. Additionally, higher liking of biology as a school subject was associated with the lower accuracy, whereas a higher interest in environment with higher accuracy.

4: The young citizen scientists provided reliable data with the exception of overrepresented false positive observations. My results suggest that the quality and quantity of citizen-generated data are not straightforwardly dependent on the selected target groups. Citizen science activities should be planned by careful consideration of the context, for example the school strongly shapes the participatory activities.

## Introduction

Citizen science is increasingly used to generate data and knowledge of the natural world (Bonney, Cooper, et al., 2009). Indeed, ecology is one of the foremost fields that has been using citizen-generated data and knowledge for a long time (Dickinson et al., 2012; Pocock, Tweddle, Savage, Robinson, & Roy, 2017). Much of the species occurrence data is collected by citizen observations, ranging from experienced amateurs, such as bird enthusiasts, to less experienced citizens providing photos on the social media (Miller-Rushing, Primack, & Bonney, 2012; McKinley et al., 2017). More broadly, citizen science is seen as a crucial step to more democratic and inclusive scientific approach (Irwin, 1995; Strasser, Baudry, Mahr, Sanchez, & Tancoigne, 2019; de Sherbinin et al., 2021)

From an ecological science point of view, the citizen science has been discussed mainly as a data generation process, and thus, the quality of the citizen science data and knowledge has been addressed in many studies (Dickinson, Zuckerberg, & Bonter, 2010; Isaac, van Strien, August, de Zeeuw, & Roy, 2014; Kelling et al., 2015; Lukyanenko, Parsons, & Wiersma, 2016; Johnston, Matechou, & Dennis, 2022). Subsequently, studies have involved validating the data with another dataset (Matutini, Baudry, Pain, Sineau, & Pithon, 2021) or by comparing citizen participants’ assessment with professional assessments (Kelling et al., 2015; Di Cecco et al., 2021). In general, the quality of the citizen scientist collected data can be improved prior to the data collection by targeting and choosing the participants or through training of the participants, while after data collection, the known biases can be taken into account during the analysis (Johnston et al., 2022).

While the effects of participation in the citizen science projects on participants’ attitudes have been previously studied (Brossard, Lewenstein, & Bonney, 2005; Bonney, Phillips, Ballard, & Enck, 2016; Kelemen-Finan, Scheuch, & Winter, 2018), it is less often studied how the attitudes of the participants shape their participation and eventually the quality of the collected data. In this study, I leveraged the setting of citizen science project, where lower (13-16-years-old) and upper (16-19-years-old) secondary school students^1^ participate in the data collection on urban rat presence or absence with easy-to-use track plates. I studied how participant characteristics relate to the quality of the collected data. This approach is crucial in understanding how scientists should choose their target groups when planning citizen science projects.

### Participants in ecological citizen science

The profiles of participants in ecological citizen science has been widely studied (Pocock et al., 2017). The current composition of white and middle-class participation echoes the past participation by the gentlemen of independent means in the scientific progress (Silvertown, 2009; Pandya & Dibner, 2018). Thus, there have been calls to involve underrepresented groups such as racial or ethnic minorities, young or older citizens and less educated or less wealthy citizens (Cooper et al., 2021; Pateman, Dyke, & West, 2021) and calls to conceptually broaden the participation to younger people and non-human animals or to pay attention to issues of epistemic pluralism such as Indigenous and Local Knowledge (Tengö, Austin, Danielsen, & Fernández-Llamazares, 2021; Rautio et al.,2022)

The central reasons for citizen participation vary from citizen science project to project. While the number of studies on the motivations of the participants has increased, they are difficult to compare as they are individual cases studies (see Jeanmougin, Levontin, & Schwartz, 2017). Jeanmougin et al., (2017) developed a framework for motivations that includes 15 different motivation categories from stimulation and hedonism to a desire for belongingness and helping. In my project, it is up for debate whether it counts as authentic citizen science as students cannot voluntarily decide their participation but rather the decision has been made by their teacher (though “Because someone asked/told me to” is covered as a motivation in West, Dyke, & Pateman, 2021). Nevertheless, the students also approach school-situated citizen science with a multitude of attitudes and motivations (Kelemen-Finan et al., 2018). These attitudes are not only important for the students’ experience of participation but also for the quality of the data collection.

### Citizen scientists’ attitudes

In this study, the most relevant participants’ attitudes are their attitude towards animals, towards biology as a school subject and towards learning about environmental issues and additionally their sensitivity to disgust. An attitude is a disposition towards a particular concrete or abstract object, person, thing or event with favour or disfavour (Eagly & Chaiken, 1993). The attitudes can be single or multiple constructs: for example, attitudes towards science learning consists of numerous subconstructs. The attitudes are expected to lead to motivation and behaviours. In contrast, interest is more specific relationship between an individual and an object, which can be situational in responses to something interesting or integral as being something more stable and gradually developing (Hidi & Renninger, 2006; Krapp, 2007).

Previous research has shown that in animal-related citizen science projects, the interest or affection towards the target species is one of the factors that drives participation and motivation (Land-Zandstra, Agnello, & Selman Gültekin, 2021). For example, ‘charismatic megafauna’ are conjured in conservation contexts to create affection between humans and wildlife (Monsarrat & Kerley, 2018). Nevertheless, some biodiversity is considered disagreeable and even disgusting, such as the focal species in my project, rats (Davey, 1994; Prokop & Tunnicliffe, 2008; Bird Rose & van Dooren, 2011). Interestingly, disgust is a rather versatile affect, which can also drive interest and transform into more positive affects and even drive learning and interest (Kolnai, n.d.; Davey, 2011; Randler, Hummel, & Wüst-Ackermann, 2013; Prokop & Fančovičová, 2017), thus this suggests that disgust-adjacent species can nevertheless stimulate participants’ interest.

As my project was carried out in the context of science education during the biology courses in lower and upper secondary school, important attitudes that relate to the participation include attitudes towards biology as a school subject and interest in learning about environmental issues. The school-situated citizen science projects have been widely studied as they are usually the most common context in which young citizens participate in citizen science. In addition to data quality, the emphasis has been strongly on the learning outcomes as the incorporation of citizen science into formal education requires considering the school curriculum (Trumbull, Bonney, Bascom, & Cabral, 2000; Cronje, Rohlinger, Crall, & Newman, 2011; Hiller & Kitsantas, 2014; Dickerson-Lange, Eitel, Dorsey, Link, & Lundquist, 2016; Shah & Martinez, 2016; Kelemen-Finan et al., 2018).

### Data generation and quality

While a large share of citizen science is driven and limited by data collection (Danielsen et al., 2009; Dickinson et al., 2012; Shirk et al., 2012), it should be noted, though, that citizen science is not necessarily limited to the data collection. Citizen science can be more citizen-driven in a sense that research questions or settings can be chosen by citizens and citizens can also analyse the results (Bonney, Ballard, et al., 2009) or it can involve citizens from project design to reporting (Stevens et al., 2014). Nevertheless, most of the ecological citizen science is limited to this crowd-sourcing type of approach (Danielsen et al., 2009; Dickinson et al., 2012). From an ecological science point of view, the majority of the literature is concerned with the impact of the properties of citizen-generated data on ecological inference (Dickinson et al., 2010; Bonney et al., 2014; Danielsen et al., 2014; Kosmala, Wiggins, Swanson, & Simmons, 2016; Johnston et al., 2022). Indeed, citizen science projects range from highly technical to simpler projects: some projects require a specialist knowledge and long-term commitment whereas other can be easily participated in for a short period without specific training or preparation (Pocock et al., 2017). Consequently, the quality of the citizen collected data varies substantially depending on the project (Kosmala et al., 2016; Aceves-Bueno et al., 2017) and the protocol chosen by a researcher is often a limiting factor in relation to the quality and quantity of the collected data (Dickinson et al., 2010).

Indeed, there are different guidelines for researchers in how to choose and approach the target group of participating citizens to maximize data quality or optimize it in relation to data quantity. The characteristics of the participants are considered through *observer quality*, i.e., how reliable data is produced by the participating citizen (Dickinson et al., 2010; Welvaert & Caley, 2016; Horns, Adler, & Şekercioğlu, 2018). Thus far, observer quality has been only considered through technical skills either directly (Sauer, Peterjohn, & Link, 1994; Genet & Sargent, 2003) or through the proxy of age (Delaney, Sperling, Adams, & Leung, 2008). To my best knowledge, only one study has previously studied how participants’ attitudes affect the collected data quality: Crall et al. (2011) compared citizen scientists versus professionals in a simulated setting while taking into account scientific literacy and attitudes but found that these were poor predictors of data collection quality and could not be used are eligibility criteria.

### Aims and research questions

My research project provides a unique setting to understand the interplay between young citizens’ attitudes, age, gender and the quality of generated data. I have organized a broad data collection with thousands of young citizens participating in rat track plate study, all ecological data is double-checked by professional scientists and large proportion of participants have completed a background questionnaire. There were two measures of data quality that the participants have generated: first whether they have been able to collect usable data at all (i.e., successfully set plates in sensible location and sent photos to a mobile application) and second, how well their identification and quantification of rat tracks matches the expert assessment.

In detail, the research questions were:

- What is the quality of data produced by young citizen scientists as measured by
  - the successful collection and submission of data and
  - the accuracy as measured by the concurrence of rat track count between citizen and professional scientist?
- Which background variables explain the data quality of the young citizen scientists?

I explored the data quality with two different approaches. First, using the whole track plate data, I measured the success rate of the participants and compared the young citizens’ assessments of the track plates and how well it matched with the expert assessment in both presence/absence of the tracks and the number of tracks. Second, using a smaller track plate data set that could be matched to participant questionnaire data, I explored how the participants’ attitudes, gender, received support and age correlated with the success and accuracy of data collection.

I hypothesized that the higher accuracy was reached by older rather than younger students and the accuracy and success would be increased in participants who are more interested in studying biology, have more pro-environmental attitudes, who are more positive about rats and have less disgust towards rats. I would expect that similar processes would operate on both ability to collect data at all and the identification or measurement errors.

## Materials and methods

My study is situated in the wider Helsinki Urban Rat Project (HURP -https://www2.helsinki.fi/en/projects/urban-rats), an ongoing multidisciplinary research project started in 2018. HURP studies rat population dynamics in urban areas and model spatiotemporal variation in rat occurrence over the whole city of Helsinki in Southern Finland. The study organism is the brown rat (*Rattus norvegicus*), for which humans usually have strong negative attitudes (Kellert, 1985; Bjerke & Østdahl, 2004). The citizen science part of HURP involves collecting data on rat presence or absence with track plates (Hacker et al., 2016). I recruited lower and upper secondary school biology teachers through a Facebook group by selecting all 33 teachers from 20 schools who had enrolled within three hours from the original posting. Teachers were not specifically trained prior to the start of the project. The project participation for individual schools starts with the visit of the researcher bringing the necessary materials and giving a one-hour classroom lecture for teacher and students. The lecture included general introduction to urban ecology, background on rats as species, aims of the study and guidance on how to use the track plates and send collected data. Teachers were then free to organize their group’s participation as suited them best. The school year is usually divided into five periods in most of the schools in Helsinki and generally the students participated within a month of the research lecture. At the end of the each period, I sent an email to teachers asking about the data collection, mentioning any submitted data which was incomplete and asked teachers to circulate the link for the project questionnaire.

The project has run since April 2018 and the data used here has been collected until the end of 2021. The profile of the participating students was not specifically surveyed: the participating schools included both the most selective upper secondary school and those lower secondary schools that receive positive discrimination funding from the city due to having a substantial number of students from immigrant backgrounds. Lower-secondary school students were 13 to 16-year-olds, where the mode was 15-year-olds, whereas the upper-secondary school students were 16 to 19-year-olds with 17 as the mode. The participating students were from Finnish, Swedish and English language schools but the questionnaire was only in Finnish, thus excluding a few students who did not know Finnish. Teachers usually participate multiple times (1-7 times) with different student groups and did not have time to prepare much for the project; thus, the first time they participated was usually not well planned. I visited lower secondary schools 23 times and upper secondary schools 56 times. Participation was focused on the first two periods (from August to November) and the last period of the school year (April and May).

### Track plates and the success and accuracy of citizen scientists

Rat presence or absence was assessed with track plates (Figure 1). The minimum requirements for participation were that there were four plates within one study area and they were photographed daily for four days after setting the plates. When sending the data, the students counted rat tracks on the plates by dividing the plate to a 5×5 grid and observing how many grid squares had rat tracks; thus, they would submit a score between 0 and 25 for each of the four observation days. The data was submitted to the database through the EpiCollect5 mobile application (Aanensen, Huntley, Feil, Al-Own, & Spratt, 2009). In general, the students were free to decide where they set the plates and whether they worked in groups or as individuals. When working in groups, the students were asked to take responsibility for all the photos of individual plates, e.g., if there were four students in a group, each student took photos and calculated rat tracks from a single plate. I asked the students to study areas in their own near-environment where they would be interested in knowing whether there are rats and set four plates within that area to places where rats would move if they were present there. I specifically used the sentence “You have to empathize with rat’s life and think in which kind of microhabitats rats generally move” and described briefly the general considerations such as cover and food sources.

**Figure 1:**
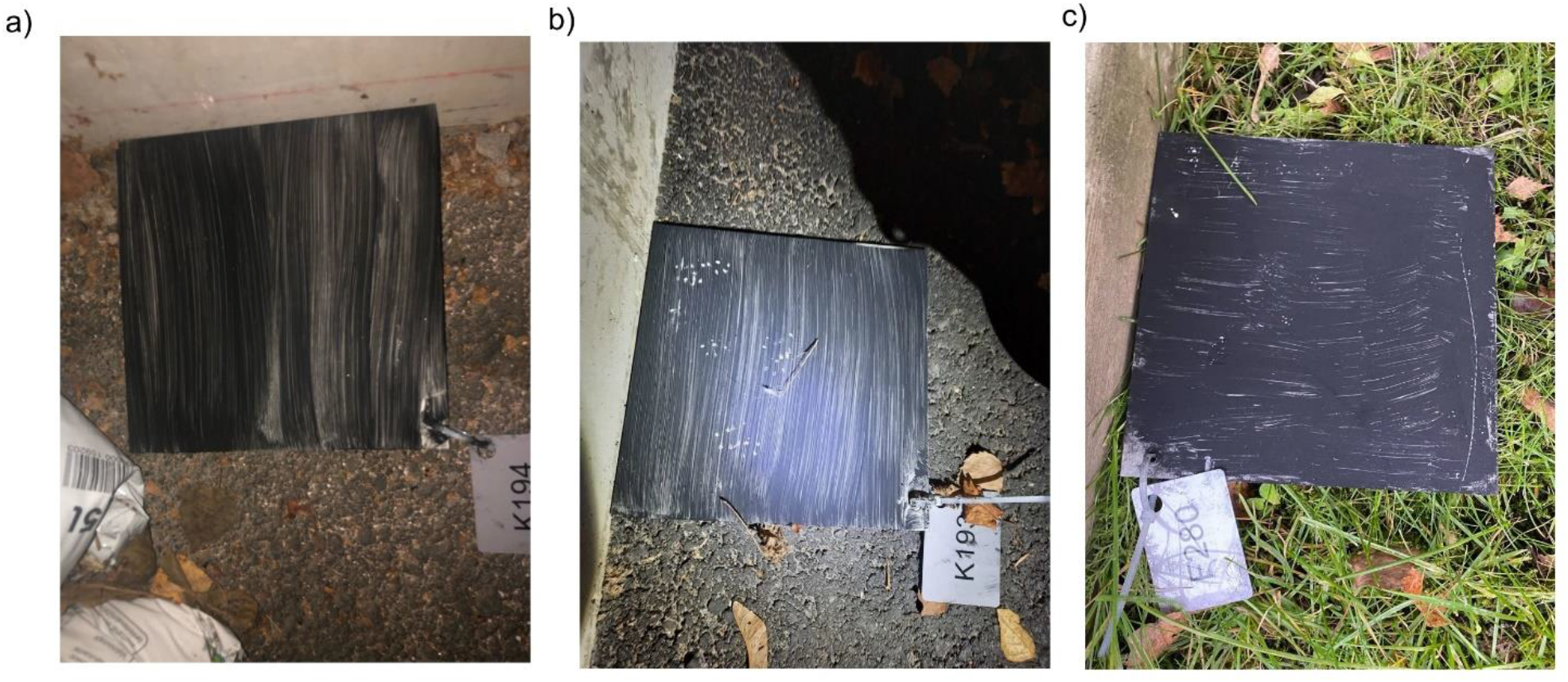
Examples of track plates a) without any tracks, b) with rat tracks and c) with tracks of smaller rodent, i.e., mice or vole.

The total number of data collection was 3,006 plates, of which 641 were collected by lower- and 2,365 by upper-secondary school students leading to an average 24 and 41 plates per visit, respectively (Table S1). After the student participation, HURP researchers checked the data integrity. The submissions that did not include photos of the plates or had missing values on location or date were deleted and considered as unsuccessful data collection. The researchers identified any tracks on the plate and noted the number of grids with rat tracks. Thus, for each plate there was a number of observed (by a young citizen scientist) and confirmed (by an experienced researcher) rat tracks. The accuracy of young ctizen scientists was measured as the congruence between the observed and confirmed tracks. The accuracy was measured as the sum of differences between observed and confirmed counts divided by the number of the plates. Thus, 0 would mean full concurrence (i.e., highest accuracy) whereas the maximum possible value (i.e., lowest accuracy) would be 100 (which corresponds to four days times 25 per day).

As a researcher, I assessed the general accuracy of the data collection by assessing the number of true and false positives and true and false negatives. True positives are plates that had confirmed rat tracks and citizen scientists had identified these as rat tracks. False positives are plates that did not have rat tracks but on which citizen scientists thought there were rat tracks. True negatives are plates which the citizen scientists correctly reported of no rat tracks and false negative are plates on which there were rat tracks, though the citizen scientists reported no tracks. This allowed me to calculate the presence-absence accuracy (true negatives and positives divided by all observations), precision (true positives divided by true and false positives), sensitivity (true positives divided by true positives and false negatives) and specificity (true negatives divided by true negatives and false positives) of presence or absence of rats in study sites.

### Questionnaire for participants

The post-participation questionnaire asked students which track plates had been used by the participants; thus, the data on the observed and confirmed number of tracks in the plates were manually added to this data. If there was a clear mention of the plates used, but they were not found in the database (i.e., they were removed during quality control) or they belonged to deleted entries, I noted these as having zero plates; therefore, they amounted to unsuccessful data collection. To link the track plate data to questionnaire data, the questionnaire had to be sent after track plate use.

To understand student attitudes for this study, four different instruments with five items per scale were combined to a questionnaire (Figure 2). I expected these previously developed and tested instruments explain how students approached the task and how reliable the data was that they were able to collect. *Liking of biology as school subject* is based on the modified version of Fennema-Sherman Mathematics Attitude Scales (Fennema & Sherman, 1976; Metsämuuronen, 2012) used in Finnish national assessments. *Interest in Environmental Issues* was based on the international ROSE questionnaire (Schreiner & Sjøberg, 2004) in which these items form a singular factor (Uitto, Juuti, Lavonen, Byman, & Meisalo, 2011). *Attitude towards Rats* was a modified scale based on an Animal Attitude Scale (Herzog, Grayson, & McCord, 2015). *Disgust* is based on the items from the revised Disgust Scale (Haidt et al., 1994, modified by Olatunji et al., 2007). These items form subscales of Core disgust and Animal-Reminder. The scales were piloted by the first two participating groups and they were deemed to perform satisfactorily.

**Figure 2:**
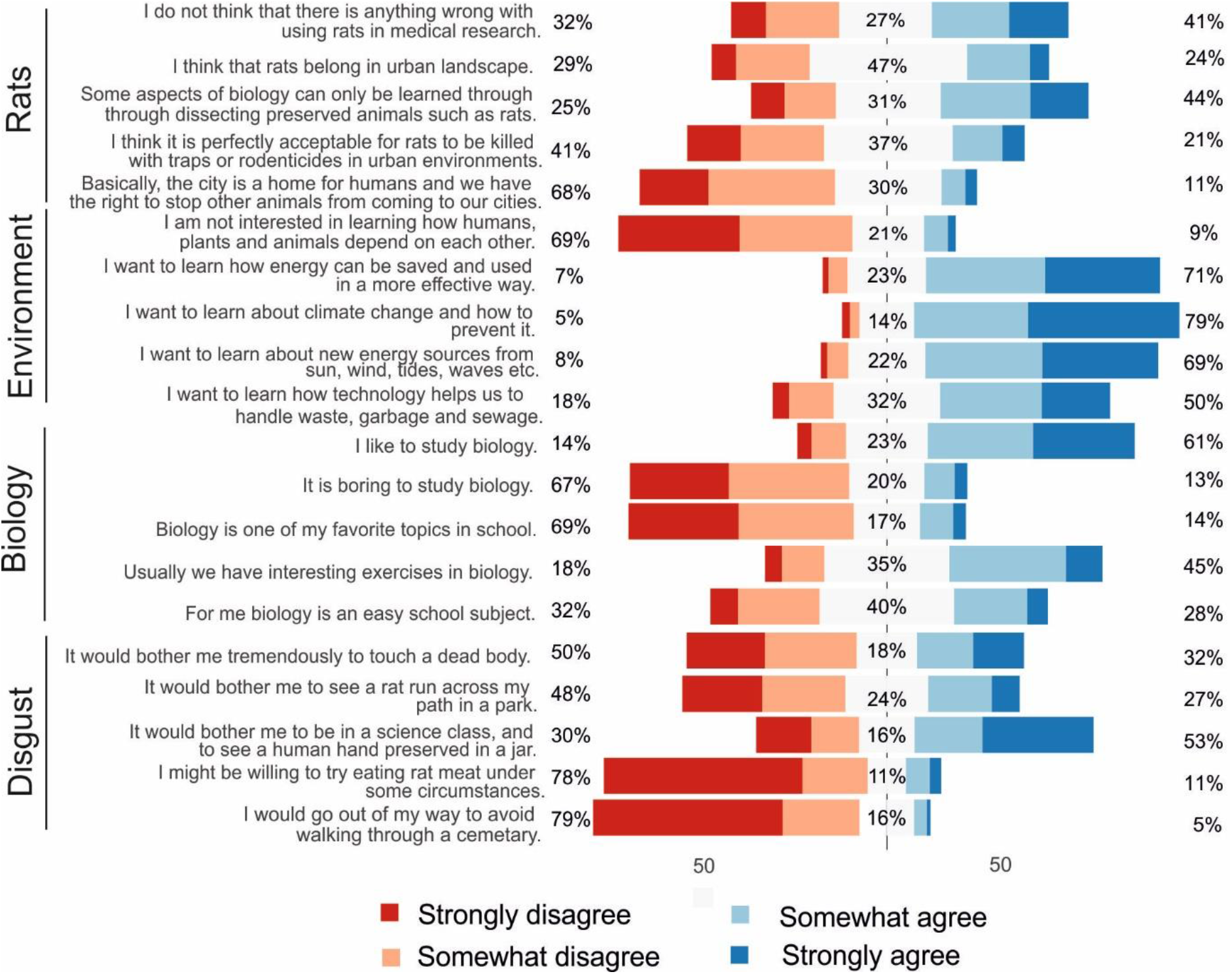
The distribution of the responses for each item grouped by scales (left side). The value on the left side indicated proportions of respondents of strongly or somewhat disagreeing, in the middle not disagreeing or agreeing and the right respondents strongly or somewhat agreeing. The item-related statistics are shown in Table S2 and S3.

The questionnaire was administered post-research: at the beginning of the project, the Epicollect5 app had a direct link to the questionnaire which participants completed out after sending data. Unfortunately, at the later stage, some of the operating systems did not work well with the link; thus in later years, the link to the questionnaire was sent to the teachers after collection of the data. In some instances, students completed the questionnaire during the classroom time, while some teachers asked students to fill this in their own time. As might be expected, this led to a lower participation rate, which could bias the results toward the more motivated students.

After collecting questionnaires, I collected further data from teachers and asked whether students received help in analyzing and counting track plate markings. I also inquired whether participation was obligatory and whether it affected the grading of the coursework. I categorized teachers’ answers in both variables to three classes.

### Statistical analysis

To calculate individual factors loadings for each of the expected attitude scales, I used item response theory (IRT). I used a generalized partial credit model (GPCM; Muraki, 1992) with Metropolis-Hastings Robbins-Monro (MH-RM; Cai, 2010a, 2010b) algorithm implemented in a mirt package (Chalmers, 2012) in R (R Core Team, 2013). MH-RM combines the flexibility of a Bayesian approach to computationally lighter maximum likelihood estimation (Cai, 2010b). GPCM is a constrained graded model that is adjusted by only a single ‘difficulty’ parameter. I assessed the usability of the factors by first doing an exploratory model, where all items were allowed to freely load on four different factors. After the exploratory model, I removed those items that loaded clearly on multiple factors and had low explanatory value (as measured by h^2^). Next, I performed a confirmatory modelling, in which I forced the items to load on predetermined factors. I assessed the model fit, item fit and personal fit. I removed the outlying respondents and redid the confirmatory analysis. Using the factor scores for each respondent, I further modelled accuracy through the covariance of four aforementioned scales.

I built generalized linear mixed models that included the school as random factor while age, gender, support, choice and factor scores for liking of biology, attitude towards rats, interest towards environmental issues and disgust were fixed factors. “Support” had three options: 1) the students did the data collection totally on their own, 2) students received help if they enquired from their teacher and 3) identification and measurement where done together in the classroom. “Choice” had also three options: 1) mandatory, 2) voluntary but students received extra credit and 3) totally voluntary. The responding variable was either the success of collecting data or the accuracy. The former included all the respondents for the questionnaire and the value was either success (1) or failure (0) using a binomial model. The latter was a numerical value between 0 and 25 and the dataset included all respondents who were successful in collecting data.

The full data set is deposited in FigShare (doi: 10.6084/m9.figshare.19583206) whereas the code for the analysis is deposited in GitHub: https://github.com/aivelo/citsci.

### Ethical considerations

Participation in the data collection for the citizen science project was part of the regular school work; therefore, the students had to participate in it with the exception of two schools were it was a voluntary part of the coursework (and students were awarded for participating). Furthermore, in three schools, the data collection included scientific report writing or other additional assignments that could affect the student grades. In relation to the questionnaire, it was made clear that participation was voluntary, participation could be ended at any time, no data would be given to their teachers and response or lack of response to the questionnaire would not affect their grades.

The first question of the questionnaire was aimed to record the informed consent of the students and 24 students did not give their consent at this stage, suggesting the students did have free choice in the questionnaire even if they had done it in the context of regular schoolwork. Nevertheless, this number of students cannot be used to calculate as the response rate as the non-consenting students could also just simply not fill the questionnaire.

Due to not handling sensitive data, there was no institutional prerequisite for an ethical review at University of Helsinki. The research permits were granted by the City of Helsinki on April 5, 2018 and September 22, 2020, respectively, and the permits from individual private schools on May 3, September 7, and October 1, 2018. All participants were informed about the aim of the study and how the materials would be collected, stored, and handled anonymously. The parents were informed beforehand in the case of students being under 15-year-olds. No personal data was collected by the HURP during the study, and the linking of the track plates and the questionnaire responses was done through track plate codes. In cases where teachers used track plates as a part of their assessment of the coursework, no information about the names was handled by HURP at any point. There was a potential risk, albeit very small, due to the amount of data, if the students collected the data from, e.g., their own home yard, that the data collection could be linked to individual data, and students were advised to not to do this, if they were concerned about it.

## Results

### The participants and their attitudes

There was a total of 772 responses to the questionnaire, of which 645 had consented to take part, completed data so that we could connect the responses to the track plate data sent through EpiCollect app and there were at least 10 responses per school. These respondents represented 1,487 plates (50% of the total plates). There was a total 12 schools (with a range of 10-224 respondents per school). The mean age of respondents was 16.56 (± 1.14); 53% were females, 43% males, 1% identified as others and 3% did not want to reveal their gender.

The respondents were generally positive towards learning biology (e.g., only 14% disagreed with the statement “I like to study biology”; Figure 2) and were very positive on learning about environmental issues (e.g., only 9% agreed with “‘I am not interested in learning how humans, animals and plants depend on each other”.) In contrast, the attitude towards rats was much more uniformly distributed, for example, for the statement “I think it is perfectly acceptable for rats to be killed with traps or rodenticides” 41% of the students disagreed and 21% agreed.

In exploratory IRT modelling, I recovered the expected factors, though not all factors performed optimally: items 9 and 20 had low factor communality values; therefore, they were dropped (Table S2). Seventeen respondents did not fit the model; they were removed from the following steps in the analysis. The confirmatory model shows an adequate fit (RMSEA = 0.062 (0.054-0.071), TLI = 0.88, CFI= 0.91; Table S2).

### The quality of data collected by young citizen scientists

Seventy-two percent of the respondents (463/645) had been successful in sending data through the mobile application which was good enough quality for the purposes of our ecological study; that meant that the plates were in an acceptable place, there was data for at least one night and the photos were clear enough to assess. The most common class was true negative occurrences, i.e., those classified as not having rat tracks by both citizen scientists and by ecologists, with 216 related responses, while true positive occurrences, i.e., classified by having rat tracks by both with 137 instances. In contrast, false positives were much more common than false negatives (78 vs. 26).

False positives relate to the tracks left by other animals, than rats, such as mice, squirrels or hedgehog which were interpreted to be rats, while false negatives were plates with rat tracks that were either near the edges of the plate or otherwise difficult to note and in a minority with the cases of other animal tracks being on the plates, suggesting that students might have not noticed rat tracks among all the other tracks.

The presence-absence accuracy, i.e., the proportion of concordant track analysis by citizen scientists and ecologists, was 77%, precision, i.e., the citizen scientists’ proportion of correctly identified presences of all presences, was 62% and sensitivity, i.e., the citizen scientists’ probability of identifying presence correctly was 82%. Finally, specificity, i.e., the citizen scientists’ probability of identifying absence correctly, was 73%.

When citizen scientists were divided to lower- and upper-secondary school groups, younger students had substantially lower values (presence-absence accuracy in upper secondary school students 79% versus 62% in lower, precision 64% vs. 59%, sensitivity 89% vs. 48%, but specificity was 73% for both). Notably, the proportion of success in collecting data differed only slightly in these groups (75% in lower-secondary school and 72% in upper-secondary school).

### The factors correlating with accuracy or overall success

Lower secondary schools had less accurate data as they had higher discrepancy between observed and confirmed values (mean 3.59 ± s.d. 10.4, vs. 1.78 ± 6.87, χ^2^_130_ = 192.3, p < 0.001). Nevertheless, the mixed-effects modelling showed that respondent age was not significant variable but rather the availability of support, rewards for voluntary participation and positive attitude towards biology as school subject were significantly positively correlated with the discrepancy between observed and confirmed values, whereas positive attitude towards learning about the environment was negatively correlated (Table 1; Figure 3). Interestingly, the situation where the whole student group identified and counted rat tracks together in classroom setting did not significantly differ from the situation where there was no support available. While the estimate of the error was substantially lower, the variation within respondents was very large.

**Table 1:**
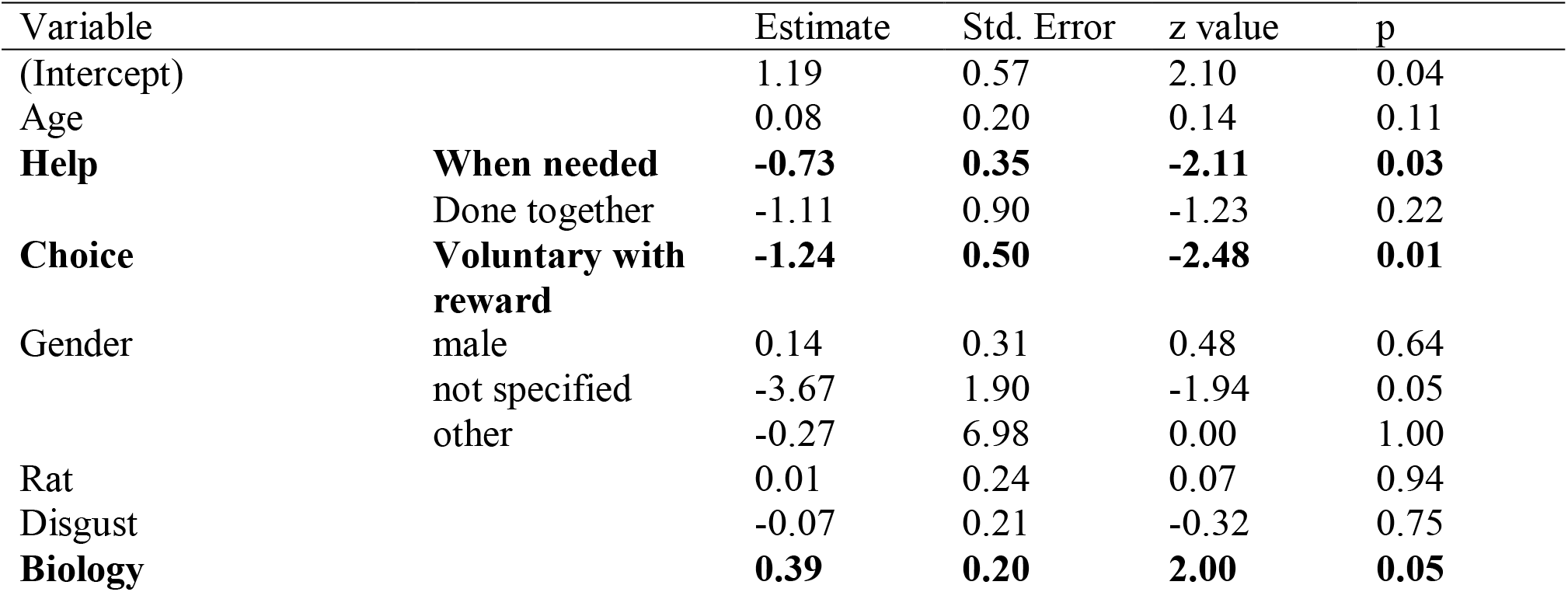
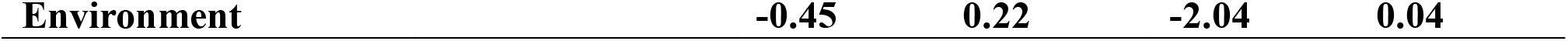
Variables affecting the accuracy. Students who received help when needed and those whose participation was voluntary but rewarded were more accurate. Additionally, those with higher liking of biology as a school subject had lower accuracy whereas those that had more interest in learning about environmental issues had higher accuracy. The baseline is female who did not receive extra help with mandatory participation.

**Figure 3:**
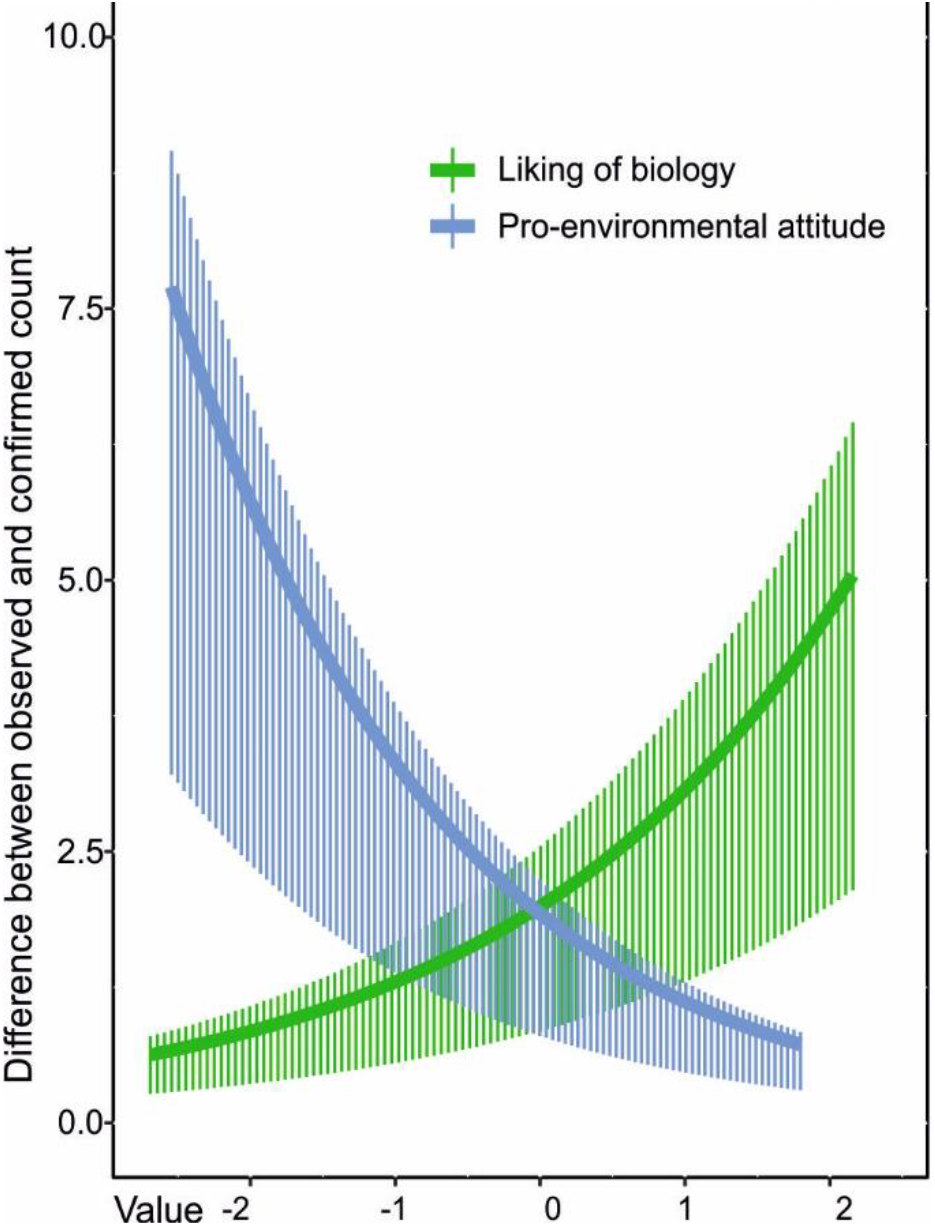
The effect sizes of the statistically significant attitude variables affecting accuracy. The X axis shows attitude values for liking of biology and pro environmental attitude. In the Y axis, the higher the difference between observed (young citizen) and confirmed (expert) measurement were, the less accurate data. Attitudes towards biology as a school subject and environmental issues were more concentrated on positive values.

When modelling accuracy by only those respondents whose plates did have confirmed rat tracks (n =163), any help or voluntary participation with rewards led to a more accurate reporting whereas men were less accurate than women (Table S5). In comparison, only rewards for voluntary participation and doing checking together correlated significantly with the likelihood of successful data collection as it lowered to success rate (Table S6). The effect of totally voluntary participation without any rewards could not be assessed in any different measures of accuracy as there was not enough cases and subsequently responses.

While the level of help, possibility of choice and age were significant variables for explaining the probability of true positives, true negatives or false negatives, none were significant for false positives (Table S4). The error rate in the plates that were linked to questionnaire responses was not statistically significant from the plates that could not be linked (i.e., questionnaire non-response) (mean 1.96 ± s.d. 7.32, vs. 1.78 ± 6.87, t_3004_= 192.3, p = 0.31).

## Discussion

My results show that young citizen scientists were comparably good in identifying the presence of rat tracks on the track plates but in contrast, they were likely to come up with false positives when there were no tracks. In general, this cannot be seen as a surprise, as the project had the explicit aim of assessing rat occurrence; thus, participants likely perceived rat tracks as an implicit goal of the study (Kervinen et al, *unpublished data*). This finding is also in line with previous research (Swanson, Kosmala, Lintott, & Packer, 2016).

Younger students (13-15-year-olds) had lower presence-absence accuracy than older students (16-19-year-olds) but there was no effect of age on measurement accuracy or overall success. The general expectation in citizen science projects involving young citizens suggests that older participants are more skilled; thus, they are expected to provide more reliable data (Delaney et al., 2008; Dickinson et al., 2010). One confusing variable which correlates with age is the organization of the school students’ participation. Other differences is that the lower-secondary school students generally participated during school time when teacher was present to help students, whereas upper-secondary school students might have completed the tasks away from school and unsupervised as homework which could have even been voluntary (Table S1). This means that lower-secondary students work more in groups and discuss the interpretation of the track plates both among their peer group and with teacher. Indeed, when included in the model, rewarded voluntary participation and available support from the teacher positively correlated with the accuracy. This might be due to selection bias of participants: those more motivated to achieve high grades are also more likely to participate and to provide quality data. In general, motivation to do “extra work” outside of the school hours might be lower without any rewards and lead to less effort in obtaining accurate data. Indeed, the total amount of plates done in upper secondary schools when participation was totally voluntary was minimal. Additionally, lower-secondary school students approach participation through group work, whereas upper-secondary students often participate alone. In general, the student performance is better in group work (Gillies, 2004; M. K. Smith et al., 2009). Nevertheless, the data here is not straightforward as both in lower- and upper-secondary school working alone led to better accuracy than working in a group (Supplementary Material). Indeed, also in cases where students worked together as a whole student group and analyzed tracks in classroom, this did not led to significantly different result than in the case where there was no support whatsoever. This is could be due to these situations making it difficult for students to opt-out and even the least motivated are compelled to participate and submit their data.

The effect of attitudes has not yet been found to affect results in citizen science projects (Crall et al., 2011; Pagès, Fischer, & van der Wal, 2018). A positive attitude towards learning biology would be expected to mean that student would try to do more careful work and follow the given guidelines more closely (San Llorente Capdevila et al., 2020). Surprisingly, in my study, those more interested in studying biology erred more often than those with less positive attitude towards biology as a school subject. Additionally, students that were more enthusiastic about studying biology were more likely to falsely detect rat tracks. The surprising correlation raises the question whether the perceived goal to find rats would be problematic especially in relation to those students who are most interested in the school subject. Then again, when lower secondary students were specifically asked whether they felt that the research was failed when they got no rat tracks, the students were able to differentiate between having a result of no rat tracks and not getting valid results at all (Kervinen & Aivelo, n.d.).

Interest in environmental attitudes correlated positively with accuracy whereas disgust or animal attitudes did not correlate at all. It is not clear why pro-environmental attitude was linked with greater accuracy. As students were prompted to think about rat presence and absence through the lens of their own near environment and to use their local knowledge, this could show a connection between learning about the environment in *general* and learning about rat presence in the near environment *specifically*. While it is difficult to assess whether studying rats was inherently more or less motivating for the students (Randler et al., 2013; Prokop & Fančovičová, 2017), at least their attitudes towards rats or disgust did not seem to play a part in the quality of their eventual data gathering. It might be also possible that in a voluntary setting, issues such as attitude towards the specific study species would be more important. Also, the variation on both attitude questions is rather small and the effect size is limited.

It should be noted that the research protocol was comparably easy as no specialist skills were needed to participate. The students were not specifically trained in identifying rat tracks, but just had a short explanation and shown photos of rat tracks and potential confounding tracks. Consequently, there were different types of negative plates: when there were no tracks whatsoever on the track plates, there is no real possibility of making mistakes. False positive cases relate to there being some tracks from other animals such as mice, squirrels or hedgehogs. Among the plates that had true occurrences of rat tracks, female students had a higher accuracy. In this case, accuracy was not so much about identifying or not identifying tracks, but rather the meticulousness of the data analysis. In general, males concentrate less on school tasks and are more overconfident (Duckworth & Seligman, 2006; Price, 2020). Interestingly, overall success of collecting any data was significantly lower in groups who had done everything together and those who had participated voluntarily. This might be a reflection on the bias that these groups had a higher proportion of unsuccessful students filling questionnaire, rather than an actual effects. Nevertheless, this suggests that the protocol was as easily followed by both lower- and upper-secondary school students.

The complex interplay among participation in citizen science, citizens’ interests and attitudes and the value of collected data requires further research. Especially in the times of a biodiversity crisis, citizen science involving living species could be used to both increase conceptual understanding of the living world and maybe create more positive relations between humans and other organisms (Aivelo, n.d.; Rautio et al., 2022). This study suggests that there are unexplored relationships between attitudes on, knowledge about and young citizens’ everyday experience of their near environment.

### Limitations

The questionnaire was submitted after the fieldwork had been completed. The underlying assumption here is that there is no substantial change in student’s attitudes during participation. In our previous work, the student interviews have shown that the students perceive that they have more positive attitudes towards rats after the research (or at least they have reflected on their relationship with rats) (Aivelo & Huovelin, 2020). Nevertheless, it is unlikely that the attitudes that are measured here change substantially during the participation (Haidt et al., 1994; Ajzen, 2001; Metsämuuronen & Tuohilampi, 2014), specifically as the participation in the research was not a planned intervention on student’s attitudes. The participation was a rather long process which ended with the track identification and counting. The questionnaires were administered right after this data collection. Thus I would argue that the questionnaires were given at a time point nearest to the step which resulted in the data quality that I have studied here.

I cannot assess how representative my sample is in relation to the all students in Helsinki (Table S1). While the sample contains schools with very different backgrounds, including language, location, student background, the actual respondents are probably biased towards the more motivated students.

There is a broad variety of ecological citizen science projects; this raises a question on how the results from my study can be generalized in relation to other citizen science projects. The research protocol was suitable for the students as there was a good success rate and there were no clear age-associated effects on how successful they were in data collection. Thus, I would expect that this study has a good generalizability to other citizen science projects. Nevertheless, it should be remembered that this study was done in a school context and school is culturally very important context for young people, which shapes all activity done within that context. Thus, I would expect that these results can be quite well generalized in participation in authentic research in a school context, but not necessarily in the voluntary participation outside of school.

My study does not yet answer to the question of how actually useful the collected data is as the modelling of the rat occurrence is yet under way. HURP has two other stakeholder-generated data sets on urban rat occurrence: trap data from an extermination company and direct observations from trash management personnel. These can be later used to validate the young citizen generated data set.

### Practical considerations for citizen science studies in schools

I have considered here only one aspect of the citizen science project data collection: the quality of the data provided by citizen scientists. In citizen science projects, citizens can also provide data that would be inaccessible to researchers otherwise. For example, in my urban rat project, students studied their own near-environments; thus, they provided local knowledge on which they were the best experts. Indeed, successful data collection further requires the imagining of rats’ experiential worlds, and I would argue that many aspects of the data collection are more than just crowdsourcing: the selection of study sites and implementation of data collection already require quite complex science skills. Thus, data quality should not be the only consideration when thinking about usability of the data but also the already existing knowledge of the participants. In ecological citizen science, this relates self-evidently for the local knowledge of participants but it can also mean that participants’ everyday experiences open new analytical potentials (Kervinen & Aivelo, n.d.).

Furthermore, it is important to remember that the ability of the participants to provide data is only one of the considerations taken in the design of citizen science studies. Important aspects include science education, democratization of science practices and knowledge creation, appreciation of indigenous or local knowledge, attitude and behaviour change, empowerment and actual change in participants’ environment (Ballard, Dixon, & Harris, 2017; Christine & Thinyane, 2021; Tengö et al., 2021; Rautio et al., 2022). These goals could and very probably will be in conflict with scientists’ interest in collecting data. For example, in a school context participation might be more useful in terms of science education for the groups who might not be as able to collect good data such as the youngest student groups. Sometimes these choices come with tradeoffs: for example, in our study, the students of the lower secondary school came generally from the vicinity of the school whereas upper secondary school students come from the area of whole city and neighboring cities. Thus, targeting lower secondary schools allowed for more targeted area of data collection.

In my study, the positive and negative cases are both valuable. In contrast, the participation was driven by a very rat-centric point of view and the students expected to find rat tracks. Thus, a limited sampling for a specific species in a given area might be expected to drive the false positive sightings but it does not seem to lead negative emotions if the species was not found (Kervinen & Aivelo, n.d.). The context of an authentic research project where the students mentioned that both presence and absence of tracks is valuable seems to partly protect from disappointments if the target species is not found.

School context strongly drives the participation as everything needs to happen within the curriculum and school year schedule. Our project was aimed at both lower- and upper-secondary school students. The much more common participation of those in upper-secondary school was likely due to difference in national curricula. The upper-secondary school curriculum includes a mandate for upper-secondary schools to both have an active cooperation with universities and to include experimental work in each biology course, whereas the lower-secondary school curriculum does not. Thus, the project provided the participating teachers a compact and easy way to fulfil those mandates. Specifically, rat tracks can be studied at any time of the year; thus there were no time limitations on our part. In general, I found enthusiastic participation in the project by students in both lower- and upper-secondary school. Students often said that they liked participation as it was “something different than everyday school” (Aivelo & Huovelin, 2020).

## Conclusion

My study shows that the quality of the collected data is not just a function of the age or skills of the participating young people, but also of how they organize their tasks and how much help they are able to get from their teacher. While student’s interest in the school subject and environmental issues correlated with the accuracy, the attitude towards the focal species or disgust sensitivity did not seem to play a part in the quality of the data.

## Acknowledgements

I thank 33 teachers and their students for participating in this research. Track plate data was also doublechecked by Suvi Sutinen, Krista Koppelomäki and Lotta Hämäläinen. I thank Álvaro Fernández-Llamazares, Aina Brias Guinart, Anttoni Kervinen, Santtu Pentikäinen and Miquel Torrents-Tico for the discussion and comments on the manuscript. Marlene Broemer did the language editing. This study was funded by Maj and Tor Nesslings Foundation, Emil Aaltonen Foundation, Kone Foundation and HiLife – Helsinki Institute of Life Sciences within the University of Helsinki.

## Footnotes

1 In this article, I use students and young citizen scientists interchangeably to acknowledge the overlapping roles of students who are participating as a part of school assignments but also as citizen scientists who are creating scientific knowledge and understanding themselves that this project was also different from the ordinary classroom assignment.

## References

Aanensen, D. M., Huntley, D. M., Feil, E. J., Al-Own, F., & Spratt, B. G. (2009). EpiCollect: Linking smartphones to web applications for epidemiology, ecology and community data collection. PLoS ONE, 4(9). doi:10.1371/journal.pone.0006968

Aceves-Bueno, E., Adeleye, A. S., Feraud, M., Huang, Y., Tao, M., Yang, Y., & Anderson, S. E. (2017). The accuracy of citizen science data: A quantitative review. Bulleting of the Ecological Society of America, 98(4), 278–290.

Aivelo, T. (n.d.). Lower and upper-secondary school students’ attitudes during citizen science project towards a representative of unloved biodiversity : urban rats. Edarxiv Preprints. doi:10.35542/osf.io/wbehu

Aivelo, T., & Huovelin, S. (2020). Combining formal education and citizen science: a case study on students’ perceptions of learning and interest in an urban rat project. Environmental Education Research, 0(0), 1–17. doi:10.1080/13504622.2020.1727860

Ajzen, I. (2001). Nature and operation of attitudes. Annual Review of Psychology, 52, 27–58.

Ballard, H. L., Dixon, C. G. H., & Harris, E. M. (2017). Youth-focused citizen science: Examining the role of environmental science learning and agency for conservation. Biological Conservation, 208, 65–75. doi:10.1016/j.biocon.2016.05.024

Bird Rose, D., & van Dooren, T. (2011). Unloved Others: Death of the Disregarded in the Time of Extinctions. Australian Humanities Review.

Bjerke, T., & Østdahl, T. (2004). Animal-related attitudes and activities in an urban population. Anthrozoos, 17(2), 109–129. doi:10.2752/089279304786991783

Bonney, R., Ballard, H., Jordan, R., McCallie, E., Phillips, T., Shirk, J., & Wilderman, C. C. (2009). Public Participation in Scientific Research: Defining the Field and Science Education. A CAISE Inquiry Group Report. Retrieved from http://www.informalscience.org/documents/PPSRreportFINAL.pdf

Bonney, R., Cooper, C. B., Dickinson, J., Kelling, S., Phillips, T., Rosenberg, K. V., & Shirk, J. (2009). Citizen science: A developing tool for expanding science knowledge and scientific literacy. BioScience, 59(11), 977–984. doi:10.1525/bio.2009.59.11.9

Bonney, R., Phillips, T. B., Ballard, H. L., & Enck, J. W. (2016). Can citizen science enhance public understanding of science? Public Understanding of Science, 25(1), 2–16. doi:10.1177/0963662515607406

Bonney, R., Shirk, J. L., Phillips, T. B., Wiggins, A., Ballard, H. L., Miller-Rushing, A. J., & Parrish, J. K. (2014). Next steps for citizen science. Science, 343(6178), 1436–1437. doi:10.1126/science.1251554

Brossard, D., Lewenstein, B., & Bonney, R. (2005). Scientific knowledge and attitude change: The impact of a citizen science project. International Journal of Science Education, 27(9), 1099–1121. doi:10.1080/09500690500069483

Cai, L. (2010a). High-dimensional exploratory item factor analysis by a Metropolis-Hastings Robbins-Monro algorithm. Psychometrika, 75(1), 33–57. doi:10.1007/s11336-009-9136-x

Cai, L. (2010b). Metropolis-Hastings Robbins-Monro algorithm for confirmatory item factor analysis. Journal of Educational and Behavioral Statistics, 35(3), 307–335. doi:10.3102/1076998609353115

Chalmers, R. P. (2012). mirt: A multidimensional item response theory package for the R environment. Journal of Statistical Software, 48(6), 1–29. doi:10.18637/jss.v048.i06

Christine, D. I., & Thinyane, M. (2021). Citizen science as a data-based practice: A consideration of data justice. Patterns, 2(4), 100224. doi:10.1016/j.patter.2021.100224

Cooper, C. B., Hawn, C. L., Larson, L. R., Parrish, J. K., Bowser, G., Cavalier, D., … Wilson, S. (2021). Inclusion in citizen science: The conundrum of rebranding. Science, 372(6549), 1386–1388. doi:10.1126/science.abi6487

Crall, A. W., Newman, G. J., Stohlgren, T. J., Holfelder, K. A., Graham, J., & Waller, D. M. (2011). Assessing citizen science data quality: An invasive species case study. Conservation Letters, 4(6), 433–442. doi:10.1111/j.1755-263X.2011.00196.x

Cronje, R., Rohlinger, S., Crall, A., & Newman, G. (2011). Does Participation in Citizen Science Improve Scientific Literacy? A Study to Compare Assessment Methods. Applied Environmental Education and Communication, 10(3), 135–145. doi:10.1080/1533015X.2011.603611

Danielsen, F., Burgess, N. D., Balmford, A., Donald, P. F., Funder, M., Jones, J. P. G., … Yonten, D. (2009). Local participation in natural resource monitoring: A characterization of approaches. Conservation Biology, 23(1), 31–42. doi:10.1111/j.1523-1739.2008.01063.x

Danielsen, F., Jensen, P. M., Burgess, N. D., Altamirano, R., Alviola, P. A., Andrianandrasana, H., … Young, R. (2014). A Multicountry Assessment of Tropical Resource Monitoring by Local Communities. BioScience, 64(3), 236–251. doi:10.1093/biosci/biu001

Davey, G. C. L. (1994). Self-reported fears to common indigenous animals in an adult UK population: The role of disgust sensitivity. British Journal of Psychology, 85(4), 541–554. doi:10.1111/j.2044-8295.1994.tb02540.x

Davey, G. C. L. (2011). Disgust: The disease-avoidance emotion and its dysfunctions. Philosophical Transactions of the Royal Society B: Biological Sciences, 366(1583), 3453–3465. doi:10.1098/rstb.2011.0039

de Sherbinin, A., Bowser, A., Chuang, T. R., Cooper, C., Danielsen, F., Edmunds, R., … Sivakumar, K. (2021). The critical importance of citizen science data. Frontiers in Climate, 3(March), 1–7. doi:10.3389/fclim.2021.650760

Delaney, D. G., Sperling, C. D., Adams, C. S., & Leung, B. (2008). Marine invasive species: Validation of citizen science and implications for national monitoring networks. Biological Invasions, 10(1), 117–128. doi:10.1007/s10530-007-9114-0

Di Cecco, G. J., Barve, V., Belitz, M. W., Stucky, B. J., Guralnick, R. P., & Hurlbert, A. H. (2021). Observing the Observers: How Participants Contribute Data to iNaturalist and Implications for Biodiversity Science. BioScience, 71(11), 1179–1188. doi:10.1093/biosci/biab093

Dickerson-Lange, S. E., Eitel, K. B., Dorsey, L., Link, T. E., & Lundquist, J. D. (2016). Challenges and successes in engaging citizen scientists to observe snow cover: From public engagement to an educational collaboration. Journal of Science Communication, 15(1), 1–14.

Dickinson, J. L., Shirk, J., Bonter, D., Bonney, R., Crain, R. L., Martin, J., … Purcell, K. (2012). The current state of citizen science as a tool for ecological research and public engagement. Frontiers in Ecology and the Environment, 10(6), 291–297. doi:10.1890/110236

Dickinson, J. L., Zuckerberg, B., & Bonter, D. N. (2010). Citizen science as an ecological research tool: Challenges and benefits. Annual Review of Ecology, Evolution, and Systematics, 41, 149–172. doi:10.1146/annurev-ecolsys-102209-144636

Duckworth, A. L., & Seligman, M. E. P. (2006). Self-discipline gives girls the edge: Gender in self-discipline, grades, and achievement test scores. Journal of Educational Psychology, 98(1), 198–208. doi:10.1037/0022-0663.98.1.198

Eagly, A. H., & Chaiken, S. (1993). The Psychology of Attitudes. Fort Worth, TX, USA: Harcourt, Brace, & Janovich.

Fennema, E., & Sherman, J. (1976). Fennema-Sherman Mathematics Attitudes Scales: Instruments designed to measure attitudes toward the learning of mathematics by females and males. Journal for Research in Mathematics Education, 7(5), 324–326.

Genet, K. S., & Sargent, L. G. (2003). Evaluation of methods and data quality from a volunteer-based amphibian call survey. Wildlife Society Bulletin, 31(3), 703–714.

Gillies, R. M. (2004). The effects of cooperative learning on junior high school students during small group learning. Learning and Instruction, 14(2), 197–213. doi:10.1016/S0959-4752(03)00068-9

Hacker, K. P., Minter, A., Begon, M., Diggle, P. J., Serrano, S., Reis, M. G., … Costa, F. (2016). A comparative assessment of track plates to quantify fine scale variations in the relative abundance of Norway rats in urban slums. Urban Ecosystems, 19(2), 561–575. doi:10.1007/s11252-015-0519-8

Haidt, J., McCauley, C., & Rozin, P. (1994). Individual differences in sensitivity to disgust: A scale sampling seven domains of disgust elicitors. Personality and Individual Differences, 16(5), 701–713. doi:10.1016/0191-8869(94)90212-7

Herzog, H. A., Grayson, S., & McCord, D. (2015). Brief measures of the animal attitude scale. Anthrozoos, 28(1), 145–152. doi:10.2752/089279315X14129350721894

Hidi, S., & Renninger, K. A. (2006). The four-phase model of interest development. Educational Psychologist, 41(2), 111–127. doi:10.1207/s15326985ep4102_4

Hiller, S. E., & Kitsantas, A. (2014). The Effect of a Horseshoe Crab Citizen Science Program on Middle School Student Science Performance and STEM Career Motivation. School Science and Mathematics, 114(6), 302–311. doi:10.1111/ssm.12081

Horns, J. J., Adler, F. R., & Şekercioğlu, Ç. H. (2018). Using opportunistic citizen science data to estimate avian population trends. Biological Conservation, 221(February), 151–159. doi:10.1016/j.biocon.2018.02.027

Irwin, A. (1995). Citizen science: a study of people, expertise and sustainable development. London, UK: Routledge.

Isaac, N. J. B., van Strien, A. J., August, T. A., de Zeeuw, M. P., & Roy, D. B. (2014). Statistics for citizen science: Extracting signals of change from noisy ecological data. Methods in Ecology and Evolution, 5(10), 1052–1060. doi:10.1111/2041-210X.12254

Jeanmougin, M., Levontin, L., & Schwartz, A. (2017). Motivations for participation to citizen-science program: A meta-analysis. Retrieved from https://cs-eu.net/sites/default/files/media/2017/07/Jeanmougin-etal-2017-STSMReportMotivationParticipation.pdf.

Johnston, A., Matechou, E., & Dennis, E. (2022). Outstanding challenges and future directions for biodiversity monitoring using citizen science data. Methods in Ecology and Evolution. doi:10.1111/2041-210X.13834

Kelemen-Finan, J., Scheuch, M., & Winter, S. (2018). Contributions from citizen science to science education: an examination of a biodiversity citizen science project with schools in Central Europe. International Journal of Science Education, 40(17), 2078–2098. doi:10.1080/09500693.2018.1520405

Kellert, S. R. (1985). American attitudes toward and knowledge of animals: An update. In M. W. Fox & L. D. Mickley (Eds.), Advances in animal welfare science 1984 (pp. 177–213). Dordrecht, Netherlands: Springer.

Kelling, S., Fink, D., La Sorte, F. A., Johnston, A., Bruns, N. E., & Hochachka, W. M. (2015). Taking a ‘Big Data’ approach to data quality in a citizen science project. Ambio, 44(4), 601–611. doi:10.1007/s13280-015-0710-4

Kervinen, A., & Aivelo, T. (n.d.). How do secondary school students handle uncertainty of authentic science inquiry? Edarxiv Preprints. doi:10.35542/osf.io/4py6n

Kolnai, A. (n.d.). Disgust. In B. Smith & C. Korsmeyer (Eds.), On disgust (pp. 29–92). Chicago, IL, USA: Open Court.

Kosmala, M., Wiggins, A., Swanson, A., & Simmons, B. (2016). Assessing data quality in citizen science. Frontiers in Ecology and the Environment, 14(10), 551–560. doi:10.1002/fee.1436

Krapp, A. (2007). An educational–psychological conceptualisation of interest. International Journal for Educational and Vocational Guidance, 7(1), 5–21. doi:10.1007/s10775-007-9113-9

Land-Zandstra, A., Agnello, G., & Selman Gültekin, Y. (2021). Participants in Citizen Science. In K. Vohland, A. Land-Zandstra, L. Ceccaroni, R. Lemmens, J. Perelló, M. Ponti, … K. Wagenknecht (Eds.), The Science of Citizen Science (p. 520). Cham, Switzerland: Springer.

Lukyanenko, R., Parsons, J., & Wiersma, Y. F. (2016). Emerging problems of data quality in citizen science. Conservation Biology, 30(3), 447–449. doi:10.1111/cobi.12706

Matutini, F., Baudry, J., Pain, G., Sineau, M., & Pithon, J. (2021). How citizen science could improve species distribution models and their independent assessment. Ecology and Evolution, 11(7), 3028–3039. doi:10.1002/ece3.7210

McKinley, D. C., Miller-Rushing, A. J., Ballard, H. L., Bonney, R., Brown, H., Cook-Patton, S. C., … Soukup, M. A. (2017). Citizen science can improve conservation science, natural resource management, and environmental protection. Biological Conservation, 208, 15–28. doi:10.1016/j.biocon.2016.05.015

Metsämuuronen, J. (2012). Challenges of the Fennema-Sherman Test in the International Comparisons. International Journal of Psychological Studies, 4(3), 1–22. doi:10.5539/ijps.v4n3p1

Metsämuuronen, J., & Tuohilampi, L. (2014). Changes in Achievement in and Attitude toward Mathematics of the Finnish Children from Grade 0 to 9—A Longitudinal Study. Journal of Educational and Developmental Psychology, 4(2), 145–169. doi:10.5539/jedp.v4n2p145

Miller-Rushing, A., Primack, R., & Bonney, R. (2012). The history of public participation in ecological research. Frontiers in Ecology and the Environment, 10(6), 285–290. doi:10.1890/110278

Monsarrat, S., & Kerley, G. I. H. (2018). Charismatic species of the past: Biases in reporting of large mammals in historical written sources. Biological Conservation, 223(April), 68–75. doi:10.1016/j.biocon.2018.04.036

Muraki, E. (1992). A Generalized Partial Credit Model: Application of an EM algorithm. Applied Psychological Measurement, 16(2), 159–176. doi:10.1177/014662169201600206

Olatunji, B. O., Williams, N. L., Tolin, D. F., Abramowitz, J. S., Sawchuk, C. N., Lohr, J. M., & Elwood, L. S. (2007). The Disgust Scale: Item Analysis, Factor Structure, and Suggestions for Refinement. Psychological Assessment, 19(3), 281–297. doi:10.1037/1040-3590.19.3.281

Pagès, M., Fischer, A., & van der Wal, R. (2018). The dynamics of volunteer motivations for engaging in the management of invasive plants: insights from a mixed-methods study on Scottish seabird islands. Journal of Environmental Planning and Management, 61(5–6), 904–923. doi:10.1080/09640568.2017.1329139

Pandya, R., & Dibner, K. A. (Eds.). (2018). Learning Through Citizen Science: Enhancing Opportunities by Design. Washington, DC, USA: National Academis of Sciences, Engineering and Medicine. doi:10.17226/25183

Pateman, R., Dyke, A., & West, S. (2021). The diversity of participants in environmental citizen science. Citizen Science: Theory and Practice, 6(1). doi:10.5334/CSTP.369

Pocock, M. J. O., Tweddle, J. C., Savage, J., Robinson, L. D., & Roy, H. E. (2017). The diversity and evolution of ecological and environmental citizen science. PLoS ONE, 12(4), 1–17. doi:10.1371/journal.pone.0172579

Price, C. R. (2020). Do women shy away from competition? Do men compete too much?: A (failed) replication. Economics Bulletin, 40(2), 1538–1547. doi:10.2139/ssrn.1444100

Prokop, P., & Fančovičová, J. (2017). The effect of hands-on activities on children’s knowledge and disgust for animals. Journal of Biological Education, 51(3), 305–314. doi:10.1080/00219266.2016.1217910

Prokop, P., & Tunnicliffe, S. D. (2008). ‘Disgusting’ animals: Primary school children’s attitudes and myths of bats and spiders. Eurasia Journal of Mathematics, Science and Technology Education, 4(2), 87–97. doi:10.12973/ejmste/75309

R Core Team. (2013). R: A language and environment for statistical computing. Vienna, Austria.: R Foundation for Statistical Computing. Retrieved from http://www.r-project.org/

Randler, C., Hummel, E., & Wüst-Ackermann, P. (2013). The Influence of Perceived Disgust on Students’ Motivation and Achievement. International Journal of Science Education, 35(17), 2839–2856. doi:10.1080/09500693.2012.654518

Rautio, P., Tammi, T., Aivelo, T., Hohti, R., Kervinen, A., & Saari, M. (2022). “For whom? By whom?”: critical perspectives of participation in ecological citizen science. Cultural Studies of Science Education, (0123456789). doi:10.1007/s11422-021-10099-9

San Llorente Capdevila, A., Kokimova, A., Sinha Ray, S., Avellán, T., Kim, J., & Kirschke, S. (2020). Success factors for citizen science projects in water quality monitoring. Science of the Total Environment, 728, 137843. doi:10.1016/j.scitotenv.2020.137843

Sauer, J. R., Peterjohn, B. G., & Link, W. A. (1994). Observer differences in the North American breeding bird survey. The Auk, 111(1), 50–62.

Schreiner, C., & Sjøberg, S. (2004). Sowing the Seeds of ROSE Background, rationale, questionnaire development and data collection for ROSE (The Relevance of Science Education) – a comparative study of students’ views of science and science education. Acta Didactica. Oslo, Norway.

Shah, H. R., & Martinez, L. R. (2016). Current Approaches in Implementing Citizen Science in the Classroom. Journal of Microbiology & Biology Education, 17(1), 17–22. doi:10.1128/jmbe.v17i1.1032

Shirk, J. L., Ballard, H. L., Wilderman, C. C., Phillips, T., Wiggins, A., Jordan, R., … Bonney, R. (2012). Public participation in scientific research: A framework for deliberate design. Ecology and Society, 17(2). doi:10.5751/ES-04705-170229

Silvertown, J. (2009). A new dawn for citizen science. Trends in Ecology & Evolution, 24(9), 467–471. doi:10.1016/j.chemosphere.2018.03.203

Smith, M. K., Wood, W. B., Adams, W. K., Wieman, C., Knight, J. K., Guild, N., & Su, T. T. (2009). Why peer discussion improves student performance on in-class concept questions. Science, 323, 122–124.

Stevens, M., Vitos, M., Altenbuchner, J., Conquest, G., Lewis, J., & Haklay, M. (2014). Taking participatory citizen science to extremes. IEEE Pervasive Computing, 13(2), 20–29. doi:10.1109/MPRV.2014.37

Strasser, B. J., Baudry, J., Mahr, D., Sanchez, G., & Tancoigne, E. (2019). ‘Citizen Science’? Rethinking science and public participation. Science & Technology Studies, 32(2), 52–76.

Swanson, A., Kosmala, M., Lintott, C., & Packer, C. (2016). A generalized approach for producing, quantifying, and validating citizen science data from wildlife images. Conservation Biology, 30(3), 520–531. doi:10.1111/cobi.12695

Tengö, M., Austin, B. J., Danielsen, F., & Fernández-Llamazares, Á. (2021). Creating synergies between citizen science and Indigenous and local Knowledge. BioScience, 71(5), 503–518. doi:10.1093/biosci/biab023

Trumbull, D. J., Bonney, R., Bascom, D., & Cabral, A. (2000). Thinking scientifically during participation in a citizen-science project. Science Education, 84(2), 265–275. doi:10.1002/(SICI)1098-237X(200003)84:2<265::AID-SCE7>3.0.CO;2-5

Uitto, A., Juuti, K., Lavonen, J., Byman, R., & Meisalo, V. (2011). Secondary school students’ interests, attitudes and values concerning school science related to environmental issues in Finland. Environmental Education Research, 17(2), 167–186. doi:10.1080/13504622.2010.522703

Welvaert, M., & Caley, P. (2016). Citizen surveillance for environmental monitoring: combining the efforts of citizen science and crowdsourcing in a quantitative data framework. SpringerPlus, 5(1). doi:10.1186/s40064-016-3583-5

West, S., Dyke, A., & Pateman, R. (2021). Variations in the motivations of environmental citizen scientists. Citizen Science: Theory and Practice, 6(1), 1–18. doi:10.5334/CSTP.370

