## Supplementary Material for "How do age, available support, gender and attitudes affect the quality of data collected by young citizen scientists in an ecological research project?"

### *Descriptive statistics of the data collection*

After data collection, each teacher described freely in an email whether the participation was voluntary or mandatory for students and whether they had additional incentives and how much help the students received in identifying and counting rat tracks. Both topics were clearly divided in three classes. In relation to the help, some student groups worked totally without any help from teacher during the identification and counting of rat tracks, while other could, if they wanted and needed, ask help from teacher. In contrast, some group identified and counted tracks all together in the classroom. The participation was mandatory in some student groups and totally voluntary in some groups. There were also groups that had voluntary participation but students received extra credit for their biology course if they participated.

**Table S1**: The number of samples per school type and the level of help students received and choice that they had on participation. Columns do not necessarily add up to the total as each teacher had consistent approach to help and choice during the study, but teacher could had different approaches within-schools and for some schools there were multiple teachers. Some teachers could teach in both lower and upper school. Total number of lower secondary schools in Helsinki is 73 and upper secondary schools 39.

| School level | Variable | | Number of schools | Number of teachers | Plates in total | Respondents in questionnaire |
|---|---|---|---|---|---|---|
| Lower | Help | No help | 2 | 2 | 192 | 8 |
| | | If asked | 2 | 2 | 60 | 0 |
| | | Done together | 4 | 5 | 389 | 54 |
| | Voluntarity | Mandatory | 8 | 9 | 641 | 62 |
| | | Extra credit | | | | |
| | | Voluntary | | | | |
| | Total | | 8 | 9 | 641 | 62 |
| Upper | Help | No help | 11 | 16 | 1438 | 260 |
| | | If asked | 4 | 7 | 751 | 251 |
| | | Done together | 2 | 3 | 176 | 72 |
| | Voluntarity | Mandatory | 12 | 21 | 1665 | 503 |
| | | Extra credit | 2 | 2 | 692 | 80 |
| | | Voluntary | 2 | 3 | 8 | 0 |
| | Total | | 14 | 26 | 2365 | 583 |
| Total | | | 20 | 33 | 3006 | 645 |

*Exploratory and confirmatory item response theory modelling*

**Table S2**: Exploratory Factor Analysis results for all the initial items. Rotated factors loadings are shown for four factors: F1 matches to "Liking of biology", F2 to "Attitude towards rats" F3 to "Disgust", and F4 to "Attitude towards learning about pro environmental choices". Communality ($h^2$) is shown for all initial items. Items in italics are inversed.

| Item | Statement | F1 | F2 | F3 | F4 | $h^2$ |
|---|---|---|---|---|---|---|
| 1 | *I do not think that there is anything wrong with using rats in medical research.* | -0.04 | -0.26 | -0.14 | -0.01 | 0.10 |
| 2 | *I am not interested in learning how humans, plants and animals depend on each other.* | -0.27 | -0.08 | -0.05 | -0.34 | 0.27 |
| 3 | I like to study biology. | 0.80 | 0.05 | 0.02 | 0.09 | 0.72 |
| 4 | It would bother me tremendously to touch a dead body. | 0.05 | 0.03 | 0.62 | -0.06 | 0.40 |
| 5 | It would bother me to see a rat run across my path in a park. | 0.02 | -0.29 | 0.35 | 0.08 | 0.18 |
| 6 | I want to learn how energy can be saved and used in a more effective way. | 0.02 | -0.10 | 0.02 | 0.80 | 0.64 |
| 7 | I think that rats belong in urban landscape. | 0.00 | 0.35 | -0.22 | 0.06 | 0.16 |
| 8 | *It is boring to study biology.* | -0.73 | -0.10 | -0.03 | -0.01 | 0.55 |
| 9 | *Some aspects of biology can only be learned by dissecting preserved animals such as rats.* | 0.01 | -0.12 | -0.17 | 0.04 | 0.05 |
| 10 | I want to learn about climate change and how to prevent it. | 0.03 | 0.13 | 0.14 | 0.66 | 0.55 |
| 11 | I think it is perfectly acceptable for rats to be killed with traps or rodenticides in urban environments. | 0.02 | 0.48 | 0.07 | -0.01 | 0.24 |
| 12 | I want to learn how new energy sources from sun, wind, tides, waves etc. | -0.02 | 0.02 | -0.13 | 0.74 | 0.55 |
| 13 | Biology is one of my favorite subjects in school. | -0.41 | 0.08 | 0.04 | 0.06 | 0.16 |
| 14 | It would bother me to be in a science class, and to see a human hand preserved in a jar. | -0.03 | 0.01 | 0.54 | 0.04 | 0.30 |
| 15 | I want to learn about technology helps us to handle waste, garbage and sewage. | -0.01 | -0.05 | -0.04 | 0.53 | 0.27 |
| 16 | Usually we have interesting exercises in biology. | 0.80 | -0.08 | -0.02 | -0.06 | 0.61 |
| 17 | *I might be willing to try eating rat meat under some circumstances.* | -0.06 | 0.05 | -0.30 | 0.04 | 0.09 |
| 18 | I would go out of my way to avoid walking through a cemetery. | 0.02 | 0.00 | 0.26 | 0.02 | 0.07 |
| 19 | For me biology is an easy school subject. | 0.54 | -0.09 | -0.12 | -0.06 | 0.31 |
| 20 | *Basically, the city is a home for humans and we have the right to stop other animals from coming to our cities.* | -0.15 | -0.10 | -0.04 | -0.09 | 0.06 |

**Table S3**: Confirmatory Factor Analysis results for all the initial items. Communality ($h^2$) and diagnostic values are shown for each items.

| Item | Statement | $h^2$ | $RMSEA_S$ $x^2$ | $p_{SC}x^2$ |
|---|---|---|---|---|
| 1 | *I do not think that there is anything wrong with using rats in medical research.* | 0.13 | 0.03 | 0.37 |
| 2 | *I am not interested in learning how humans, plants and animals depend on each other.* | 0.20 | 0.03 | 0.07 |
| 3 | I like to study biology. | 0.67 | 0.03 | 0.37 |
| 4 | It would bother me tremendously to touch a dead body. | 0.51 | 0.03 | 0.07 |
| 5 | It would bother me to see a rat run across my path in a park. | 0.04 | 0.03 | 0.07 |
| 6 | I want to learn how energy can be saved and used in a more effective way. | 0.63 | 0.03 | 0.30 |
| 7 | I think that rats belong in urban landscape. | 0.03 | 0.00 | 0.64 |
| 8 | *It is boring to study biology.* | 0.49 | 0.03 | 0.37 |
| 10 | I want to learn about climate change and how to prevent it. | 0.44 | 0.03 | 0.11 |
| 11 | I think it is perfectly acceptable for rats to be killed with traps or rodenticides in urban environments. | 0.28 | 0.03 | 0.07 |
| 12 | I want to learn how new energy sources from sun, wind, tides, waves etc. | 0.51 | 0.03 | 0.07 |
| 13 | Biology is one of my favorite subjects in school. | 0.14 | 0.03 | 0.07 |
| 14 | It would bother me to be in a science class, and to see a human hand preserved in a jar. | 0.28 | 0.03 | 0.07 |
| 15 | I want to learn about technology helps us to handle waste, garbage and sewage. | 0.26 | 0.00 | 0.57 |
| 16 | Usually we have interesting exercises in biology. | 0.61 | 0.00 | 0.69 |
| 17 | *I might be willing to try eating rat meat under some circumstances.* | 0.06 | 0.03 | 0.19 |
| 18 | I would go out of my way to avoid walking through a cemetery. | 0.07 | 0.03 | 0.37 |
| 19 | For me biology is an easy school subject. | 0.30 | 0.03 | 0.07 |

*The factors correlating with true and false positives and negatives*

**Table S4a**: The variables affecting the occurrence of true positive track plate observation. The statistically significant factors were age which means that older participants were more likely to have true positive observations as did those who assessed plates together in the classroom. The baseline is female who did not receive extra help with mandatory participation.

| Variable | | Estimate | Std. Error | z value | p |
|---|---|---|---|---|---|
| **(Intercept)** | | **-1.06** | **0.23** | **-4.62** | **<0.01** |
| **Age** | | **0.37** | **0.13** | **2.87** | **<0.01** |
| **Help** | When needed | -0.37 | 0.27 | -1.34 | 0.17 |
| | **Done together** | **1.22** | **0.39** | **3.11** | **<0.01** |
| Choice | Voluntary with extra credit | -0.38 | 0.34 | 1.10 | 0.27 |
| Gender | male | 0.22 | 0.24 | 0.90 | 0.37 |
| | not specified | 1.05 | 0.58 | 1.81 | 0.07 |
| | other | -0.26 | 3.37 | 0.02 | 0.98 |
| Rat | | 0.21 | 0.16 | 1.32 | 0.19 |
| Disgust | | -0.11 | 0.14 | -0.77 | 0.44 |
| Biology | | 0.05 | 0.14 | 0.35 | 0.73 |
| Environment | | -0.05 | 0.15 | -0.38 | 0.70 |

**Table S4b**: The variables affecting the occurrence of false positive track plate observation. No statistically significant factors found. The baseline is female who did not receive extra help with mandatory participation. When false positives were compared to only those reporting any tracks (true and false positives) or not having any tracks (false positives and true negatives), no significant variables were found.

| Variable | | Estimate | Std. Error | z value | p |
|---|---|---|---|---|---|
| **(Intercept)** | | **-1.54** | **0.26** | **-5.77** | **<0.01** |
| Age | | 0.06 | 0.17 | 0.35 | 0.73 |
| Help | When needed | 0.01 | 0.31 | 0.05 | 0.96 |
| | Done together | -0.34 | 0.52 | -0.65 | 0.52 |
| Choice | Voluntary with extra credit | -0.48 | 0.47 | -1.02 | 0.31 |
| Gender | male | 0.21 | 0.28 | 0.75 | 0.42 |
| | not specified | -0.20 | 15.3 | -0.01 | 0.99 |
| | other | -0.20 | 37.1 | -0.01 | 0.99 |
| Rat | | 0.04 | 0.19 | 0.23 | 0.82 |
| Disgust | | 0.04 | 0.17 | 0.26 | 0.80 |
| Biology | | 0.12 | 0.17 | 0.68 | 0.50 |
| Environment | | -0.20 | 0.18 | -1.12 | 0.27 |

**Table S4c**: The variables affecting the occurrence of true negative track plate observation. The statistically significant factors were voluntary participation with rewards and assessing plates together in the classroom, which both led to smaller probability of true negative plates. The baseline is female who did not receive extra help with mandatory participation.

| Variable | | Estimate | Std. Error | z value | p |
|---|---|---|---|---|---|
| (Intercept) | | 0.09 | 0.20 | 0.45 | 0.65 |
| Age | | -0.19 | 0.12 | -1.53 | 0.13 |
| **Help** | When needed | 0.32 | 0.24 | 1.34 | 0.18 |
| | **Done together** | **-1.13** | **0.39** | **-2.87** | **<0.01** |
| **Choice** | **Voluntary with extra credit** | **-0.71** | **0.34** | **-2.05** | **0.04** |
| Gender | male | -0.24 | 0.22 | -1.19 | 0.24 |
| | not specified | -0.32 | 0.57 | -0.55 | 0.57 |
| | other | 3.26 | 71.8 | 0.01 | 0.99 |
| Rat | | -0.09 | 0.15 | -0.60 | 0.55 |
| Disgust | | 0.06 | 0.13 | 0.49 | 0.62 |
| Biology | | -0.06 | 0.13 | -0.46 | 0.65 |
| Environment | | 0.08 | 0.14 | 0.58 | 0.59 |

**Table S4d**: The variables affecting the occurrence of false negative track plate observation. The only statistically significant factor was voluntary participation with rewards, which smaller probability of having false negative observation. The baseline is female who did not receive extra help with mandatory participation.

| Variable | | Estimate | Std. Error | z value | p |
|---|---|---|---|---|---|
| **(Intercept)** | | **-3.90** | **0.76** | **-5.15** | **<0.01** |
| Age | | -0.69 | 0.39 | -1.76 | 0.08 |
| Help | When needed | -0.27 | 1.10 | -0.24 | 0.81 |
| | Done together | 0.39 | 1.06 | 0.36 | 0.71 |
| **Choice** | **Voluntary with extra credit** | **-2.99** | **1.03** | **2.93** | **<0.01** |
| Gender | male | -0.58 | 0.59 | -0.96 | 0.32 |
| | not specified | -0.63 | 1.27 | -0.49 | 0.62 |
| | other | -2.01 | 72.4 | -0.03 | 0.98 |
| Rat | | -0.62 | 0.40 | -1.54 | 0.12 |
| Disgust | | -0.04 | 0.33 | -0.12 | 0.91 |
| Biology | | -0.33 | 0.33 | -1.02 | 0.31 |
| Environment | | 0.59 | 0.36 | 1.64 | 0.10 |

*The factors correlating with accuracy or success*

**Table S5:** The variables affecting the accuracy when only considering those track plates which had rat tracks based on the expert assessment. The statistically significant factors were the male gender which had substantially lower accuracy, voluntary participation with rewards and either having help when needed or assessing plates together in the classroom, which led to higher accuracy. The baseline is female who did not receive extra help with mandatory participation.

| Variable | | Estimate | Std. Error | z value | p |
|---|---|---|---|---|---|
| **(Intercept)** | | **-1.04** | **0.35** | **-2.93** | **<0.01** |
| Age | | -0.04 | 0.21 | -0.20 | 0.84 |
| **Help** | **When needed** | **-0.93** | **0.39** | **-2.36** | **0.02** |
| | **Done together** | **-1.62** | **0.71** | **-2.30** | **0.02** |
| **Choice** | **Voluntary with extra credit** | **-0.95** | **0.48** | **-1.99** | **0.05** |
| **Gender** | **male** | **1.01** | **0.36** | **2.78** | **<0.01** |
| | not specified | -0.55 | 1.18 | -0.47 | 0.64 |
| Rat | | 0.22 | 0.28 | 0.78 | 0.44 |
| Disgust | | 0.30 | 0.24 | 1.23 | 0.22 |
| Biology | | 0.30 | 0.19 | 1.54 | 0.12 |
| Environment | | -0.15 | 0.25 | -0.58 | 0.56 |

**Table S6:** The variables affecting the success of the data collection. The significant variables were voluntary participation with rewards and when the track plate checking was done together, which both decreased success rate. The baseline is female who did not receive extra help with mandatory participation.

| Variable | | Estimate | Std. Error | z value | p |
|---|---|---|---|---|---|
| (Intercept) | | 0.71 | 0.34 | 2.06 | <0.01 |
| Age | | 0.09 | 0.18 | 0.49 | 0.75 |
| **Help** | When needed | -0.28 | 0.64 | -0.43 | 0.66 |
| | **Done together** | **-0.22** | **0.64** | **-2.61** | **<0.01** |
| **Choice** | **Voluntary with extra credit** | **-1.25** | **0.47** | **-2.32** | **0.02** |
| Gender | male | -0.18 | 0.23 | -0.80 | 0.43 |
| | not specified | -0.39 | 0.58 | -0.67 | 0.51 |
| | other | -0.27 | 1.32 | -0.67 | 0.84 |
| Rat | | -0.08 | 0.15 | -0.51 | 0.61 |
| Disgust | | 0.05 | 0.14 | 0.37 | 0.71 |
| Biology | | 0.02 | 0.13 | 0.20 | 0.84 |
| Environment | | 0.20 | 0.14 | 1.48 | 0.14 |

*The effect of single or group work on the reliability*

Anecdotally, the lower secondary school students worked more often in groups. To explore whether this affected the reliability of their results, I looked at the correlations and interactions between error rates, lower or upper school level and whether the students worked alone or in group. The data is not optimal for this analysis, as I do not have direct data on students group work. I assessed this indirectly so that any students who reported less than four plates, where inferred to work with other students, as the four plates was the standard number of plates on a site. It was more difficult to assess where the cases where all students working in the same group reported all four plates. I identified these cases in the data by looking at the reports of overlapping plate numbers in same time frames. This is prone to errors, though, as not all students answered to the questionnaire and thus some students, which I assessed as working alone where in fact working in group.

To assess the effects of school level and mode of working, I built generalized linear mixed models, which included the school as random factor and school level (lower/upper) and the mode of working (individual/group) as fixed factors with interaction. No variable was statistically significant: those participating alone had lower error rate (estimate: -0.92, standard error 0.55, z = -1.6, p = 0.09), the error rate was higher in upper secondary school (0.33, SE 0.39, z =  0.8, p = 0.40) and there was a positive interaction between working alone and upper secondary school (0.64, SE 0.633, z = 1.0, p = 0.31).

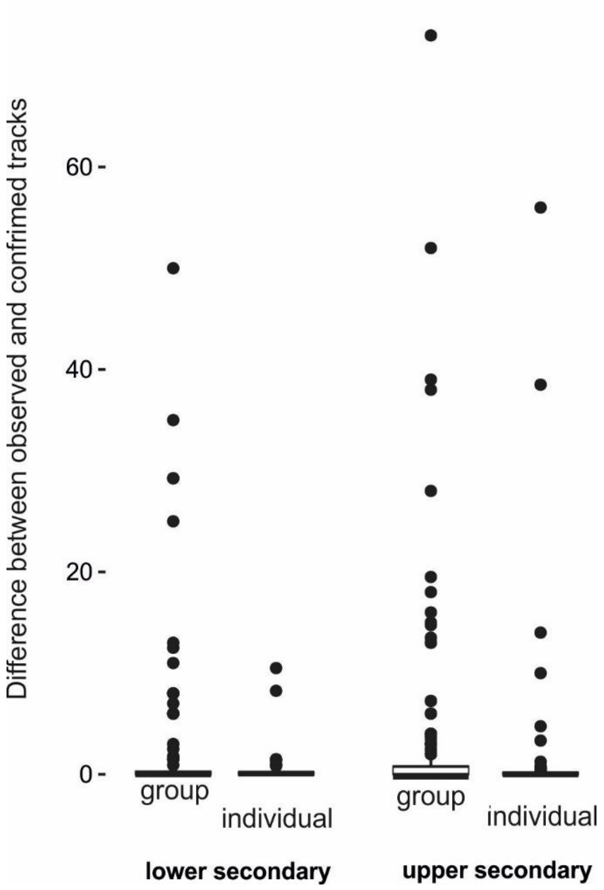


**Figure S1:** The error rate in lower and upper secondary group students depending whether they worked alone or as a group.
